# Biased Auditory Nerve Central Synaptopathy Exacerbates Age-related Hearing Loss

**DOI:** 10.1101/2020.06.09.142737

**Authors:** Meijian Wang, Chuangeng Zhang, Shengyin Lin, Yong Wang, Benjamin J. Seicol, Robert W. Ariss, Ruili Xie

**Affiliations:** Department of Otolaryngology – Head and Neck Surgery, The Ohio State University, Columbus, OH 43210, USA; College of Medicine and Life Sciences, University of Toledo, Toledo, OH 43614, USA; Department of Neuroscience, The Ohio State University, Columbus, OH 43210, USA

**Keywords:** Auditory nerve central synapse, synaptopathy, age-related hearing loss, spiral ganglion neuron, calretinin, VGluT1, endbulb of Held, bushy neuron, cochlear nucleus, synaptic convergence

## Abstract

Sound information is transmitted from the cochlea to the brain by different subtypes of spiral ganglion neurons (SGN), which show varying degrees of vulnerbility under pathological conditions. It remains unclear how information from these SGNs reassemble among target neurons in the cochlear nucleus (CN) at the auditory nerve (AN) central synapses, and how different synapses change during hearing loss. Combining immunohistochemistry with electrophysiology, we investigated the giant endbulb of Held synapses and their postsynaptic bushy neurons in mice under normal hearing and age-related hearing loss (ARHL). We found that calretinin-expressing and non-calretinin-expressing endbulbs converge at continuously different ratios onto bushy neurons with varying physiological properties. Endbulbs degenerate during ARHL, and the degeneration is more severe in non-calretinin-expressing synapses, which correlates with a gradual decrease in neuronal subpopulation predominantly innervated by these inputs. Our findings suggest that biased AN central synaptopathy and shifted CN neuronal composition underlie reduced auditory input and altered central auditory processing during ARHL.

## INTRODUCTION

Spiral ganglion neurons (SGN) of the peripheral auditory system convey sound information from sensory hair cells to the cochlear nucleus (CN) (Nayagam et al., 2011), which is the first station of the central auditory system. Most SGNs (90-95%) are type I (Kiang et al., 1982; Spoendlin, 1969) and can be divided into three subtypes based on their spontaneous firing rate and threshold, fiber caliber and preferential terminal distribution around the hair cell basal circumference (Liberman, 1978, 1980, 1982a, b; Liberman and Oliver, 1984). More recently, these subtypes of type I SGNs were shown to have distinct molecular signatures, including calretinin as a marker for type I_a_ SGNs with high spontenous rate/low threshold (Petitpre et al., 2018; Shrestha et al., 2018; Sun et al., 2018). These SGN subtypes differentially encode sound intensity, and collectively ensure a comprehensive representation of the acoustic environment. Transfer of sound information through the bipolar SGNs begins at their dendritic terminal of cochlear synapses on hair cells, and ends at their central auditory nerve (AN) synapses in the CN (Nayagam et al., 2011; Yu and Goodrich, 2014). Pathology of the cochlear synapses (known as “cochlear synaptopathy”) from hearing loss and aging preferentially affects the low spontaneous rate/high threshold subtype of type I SGNs (Furman et al., 2013; Liberman et al., 2015), and is widely recognized as a major mechanism that underlies hidden hearing loss (Kujawa and Liberman, 2015; Liberman et al., 2015; Liberman, 2017). However, it is not known how different subtypes of AN synapses converge onto individual CN neurons (Nayagam et al., 2011; Spirou et al., 2005; Yu and Goodrich, 2014), how different innervation patterns correlate with properties of postsynaptic CN neurons, and most importantly, how subtype-specific changes of AN synapses can affect neural processing by target CN neurons under pathological conditions.

The most prominent AN synapse is the large endbulb of Held (Rouiller et al., 1986; Ryugo and Fekete, 1982), which reliably transmits information about the temporal fine structure (TFS) of sound to its postsynaptic bushy neuron in ventral CN (Joris et al., 1994a; Joris et al., 1994b; Manis et al., 2011). Such TFS information is crucial for pitch perception and speech recognition (Moore, 2008; Shannon et al., 1995), which are often compromised in hearing impaired patients (Anderson et al., 2012; Grose and Mamo, 2010; Lorenzi et al., 2006). After hearing loss, endbulb of Held synapses show modified synaptic morphology with reduced size and simplified complexity (Ryugo et al., 1997; Ryugo et al., 1998; Wright et al., 2014), as well as altered function associated with compromised synaptic transmission (Oleskevich and Walmsley, 2002; Wang and Manis, 2005; Wright et al., 2014; Xie and Manis, 2017b; Zhuang et al., 2017), which degrades the firing of postsynaptic bushy neurons (Xie, 2016). Despite decades of research on the endbulbs of Held, little is known about the convergence of—as well as how synaptopathy occurs among—different subtypes of endbulb of Held synapses during hearing loss. Furthermore, the impacts of subtype-specific synaptopathy on the response properties of postsynaptic bushy neurons remain unclear.

To investigate these questions, we combined immunohistochemistry with electrophysiology using acute brain slices from CBA/CaJ mice at three age groups: young mice (1.5-4.5 months) with normal hearing, middle-aged mice (17-19 months) with moderate ARHL that mimics hidden hearing loss (Sergeyenko et al., 2013), and old mice (28-32 months) with prominent ARHL. Endbulb synapses from type I_a_ SGNs (high spontaneous rate/low threshold) were differentiated from the other subtypes (type I_b_ or I_c_ SGNs with medium/low spontaneous rate and medium/high threshold) based on immunostaining for calretinin (Petitpre et al., 2018; Shrestha et al., 2018; Sun et al., 2018). We report the innervation patterns of type I_a_ and non-type I_a_ endbulbs on individual bushy neurons and their intrinsic and AN-evoked response properties, as well as subtype-specific AN central synaptopathy during ARHL.

## RESULTS

### Multiple subtypes of AN central synapses converge onto individual CN neurons

We performed current clamp recording from bushy neurons in parasagittal CN slices and filled the target neurons with Alexa Fluor 594, which was included in the electrode solution (Fig. 1A-B). Slices were post hoc immunostained using antibodies against VGluT1, which labels glutamate vesicles in all AN synaptic terminals, and calretinin, which labels only the AN fibers and synapses from type I_a_ SGNs (STAR methods). As shown in Fig. 1C, the bushy neuron receives inputs from type I_a_ endbulb of Held synapses (red), which contains glutamate vesicles revealed as VGluT1-labeled puncta (yellow in merged panel). In addition, there are VGluT1-labeled puncta (green in merged panel) that are not enclosed in calretinin-labeled synapses, indicating that they are from non-type I_a_ AN synapses (presumably type I_b_ or I_c_). Z-stack images were used to reconstruct the 3-dimentional structure of the type I_a_ endbulbs as well as VGluT1-labeled puncta surrounding the soma of the labeled bushy neuron (Fig. 1D-H; Supplemental Video 1). This approach reveals that type I_a_ and non-type I_a_ AN synapses converge onto individual CN neurons (Fig. 1F).

**Figure 1.**
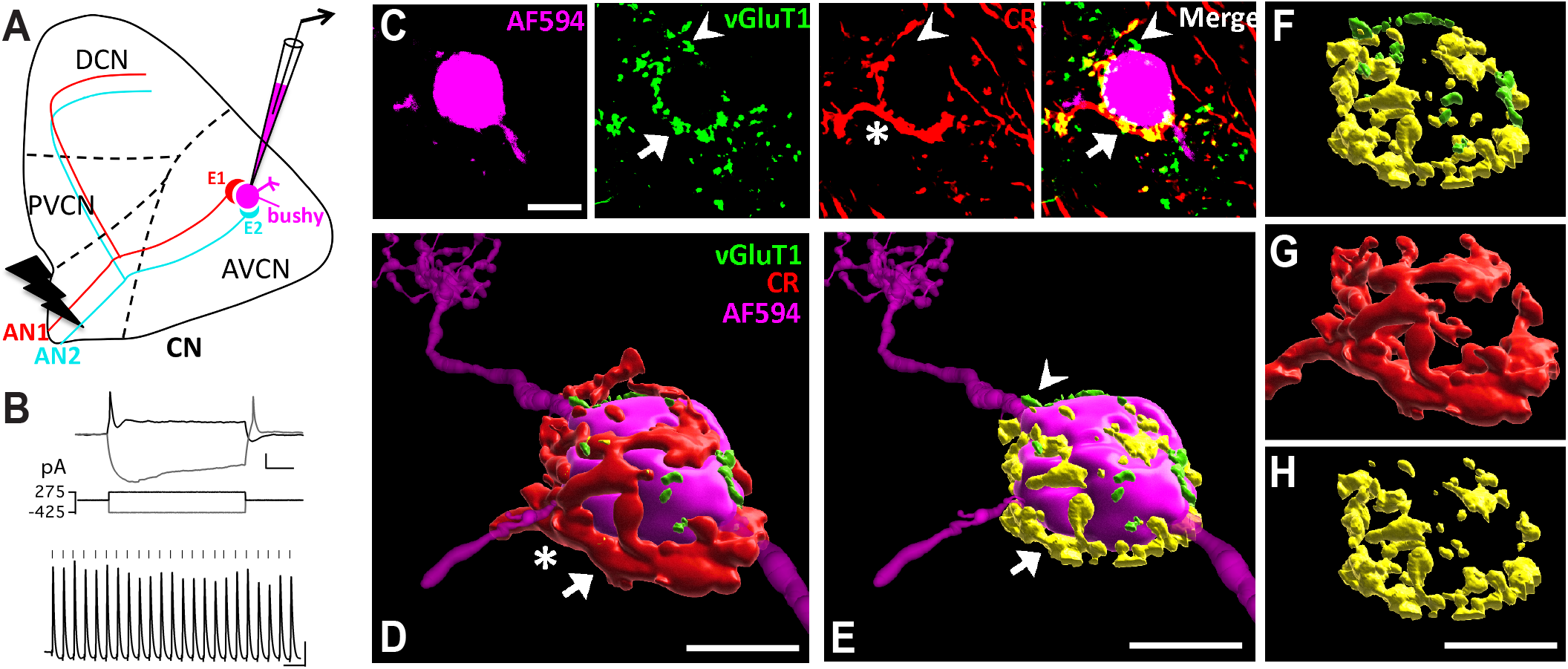
Multiple subtypes of AN synapses converge onto individual CN neurons. (**A**) Diagram of the CN and experimental setup. It depicts two endbulb of Held synapses (E1 and E2) from different AN fibers (AN1 and AN2) that innervate the target bushy neuron, which is recorded and filled with fluorescent dye (shown in magenta). AN is activated by electric stimulation. CN: cochlear nucleus; AVCN: anteroventral CN; PVCN: posteroventral CN; DCN: dorsal CN. (**B**) Example responses of a bushy neuron to current step injections (top) and trains of stimulation at AN (bottom). Ticks: stimulus onset. Scale: 10 mV and 20 ms. (**C**) Single frame confocal images of the bushy neuron in (**B**) filled with Alexa Fluor 594 dye (magenta) during whole-cell patch clamp recording. The slice was then immunostained against VGluT1 (green) and calretinin (CR; red). A single type I_a_ endbulb of Held synapse is labeled by both calretinin (star) and VGluT1 (arrow). The overlap-labeled puncta are shown in yellow in merged panel (arrow). VGluT1-labeled puncta that do not overlap with CR staining are shown in green (arrowhead), which are from AN synapses that do not express calretinin (non-type I_a_ synapse). Scale: 10 μm. (**D**) 3D reconstruction of the bushy neuron and AN synapses in (**C**). Red: CR-stained type I_a_ endbulb of Held synapse. Scale: 10 μm, also applies to **E-H**. See also Supplemental Video 1. (**E**) same as in (**D**) except revealing only the VGluT1-labeled puncta. Yellow (arrow): VGluT1-labeled puncta inside the calretinin-expressing type I_a_ synapses; green (arrowhead): puncta from non-calretinin-expressing (non-type I_a_) synapses. (**F**) View of all VGluT1-labeled puncta from (**E**). Note that this bushy neuron received mostly type I_a_ synaptic inputs from the AN. (**G**) Morphology of a single type I_a_ endbulb of Held synapse from (**D**). (**H**) VGluT1-stained puncta inside the single endbulb in (**G**).

### The proportion of type I_a_ synaptic inputs correlates with physiological properties of postsynaptic neurons

We studied the innervation pattern of type I_a_ and non-type I_a_ synapses onto 49 bushy neurons and their response properties using young mice (Fig. 2A-L). The ratio of convergence between different synaptic inputs varied on a continuum among bushy neurons from type I_a_-dominant (I_a_-D; with I_a_ volume > 50%) to non-type I_a_-dominant (Non-I_a_-D; with I_a_ volume < 50%), as quantified by the volume of VGluT1-labeled puncta from each subtype (Fig. 2M). In addition, the proportion of type I_a_ inputs onto a postsynaptic bushy neuron correlates with its intrinsic properties and AN-evoked firing properties (Fig. 2A-L, N-O). Compared to Non-I_a_-D neurons, I_a_-D neurons showed significantly more depolarized resting membrane potential (Fig. 2P), smaller spike amplitude (Fig. 2R), required less current injection to trigger threshold spikes (Fig. 2S), but had similar input resistance (Fig. 2Q). Therefore, bushy neurons that receive more type I_a_-inputs (I_a_-D) are more excitable than those that receive more non-type I_a_ inputs (Non-I_a_-D). Consistently, trains of AN stimulation at 100 Hz evoked sustained spikes with moderate adaptation in I_a_-D neurons (Fig. 2C, F), but only elicited transient or onset spikes in Non-I_a_-D neurons (Fig. 2I, L). In summary, neurons receiving mostly type I_a_ inputs showed significantly higher AN-evoked firing rate (Fig. 2T) and less spike adaptation (Fig. 2U). They also fired spikes with higher temporal precision (Fig. 2V). Similar differences were also observed in the responses to trains of AN stimulation at 400 Hz (Fig. S1). These results indicate that AN synapses from different subtypes of SGNs are not uniformly distributed onto populations of CN bushy neurons, and that the pattern of synaptic input is correlated with the intrinsic excitability of target cells.

**Figure 2.**
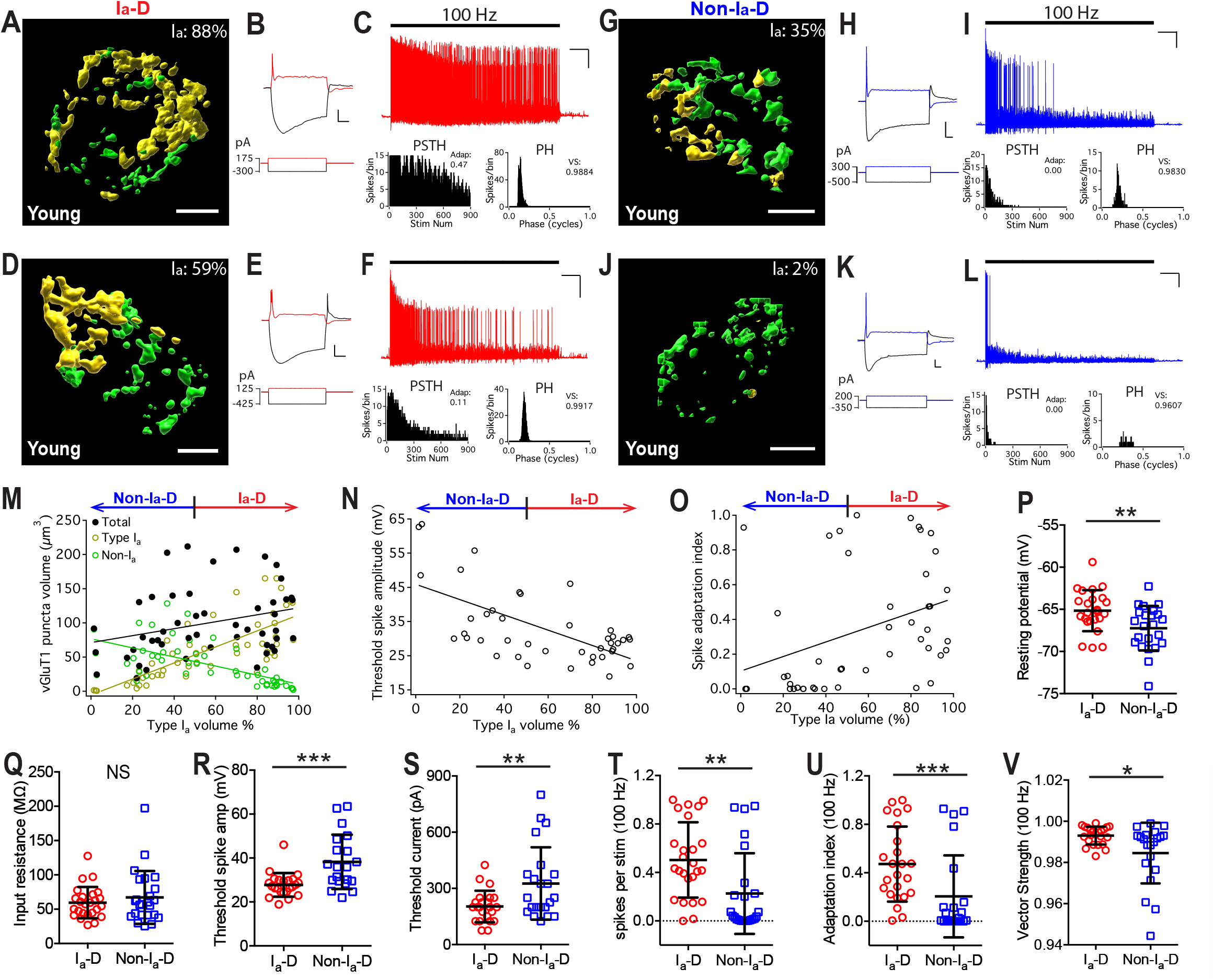
The proportion of type I_a_ synaptic inputs correlates with physiological properties of postsynaptic neurons. (**A-C**) Example bushy neuron that received mostly type I_a_ synaptic inputs (I_a_-dominant, or I_a_-D) from the AN. (**A**) Reconstructed VGluT1-stained puncta surrounding the soma of the bushy cell (not shown) from type I_a_ synapses (yellow) and non-type I_a_ synapses (green). I_a_%: volume percentage of the VGluT1-stained puncta from type I_a_ synapses over total. Scale: 5 μm. (**B**) Responses of the bushy cell to current step injections. Red traces: threshold current injection (bottom) and response (top). Scale: 10 mV and 20 ms. (**C**) The neuron fired sustained spikes to a train of auditory nerve stimulation (black bar) at 100 Hz. Scale: 10 mV and 1 s. PSTH: post-stimulus time histogram. Adap: spike adaptation index. PH: period histogram. VS: vector strength. (**D-F**) Example bushy neuron with I_a_-D but lower proportion of type I_a_ inputs. (**G-I**, **J-L**) Two example bushy neurons that received mostly non-type I_a_ synaptic inputs (Non-type I_a_-dominant, or Non-I_a_-D), and fired only transient or onset spikes to AN stimulus trains. (**M**) Bushy neurons with different fraction of type I_a_ inputs (x-axis) show correlated distribution in their total volume of VGluT1-stained puncta (black), volume of type I_a_ only puncta (yellow), and volume of non-type I_a_ only puncta (green). Linear regression lines: black, r^2^ = 0.09, p = 0.039; yellow, r^2^ = 0.57, p < 0.0001; green, r^2^ = 0.38, p < 0.0001. (**N**) Bushy neurons with different fraction of type I_a_ inputs (x-axis) show correlated difference in threshold spike amplitude, as shown in **B**, **E**, **H** and **K**. Linear regression line: r^2^ = 0.40, p < 0.0001. (**O**) Bushy neurons with different fraction of type I_a_ inputs (x-axis) show different firing patterns to 100 Hz stimulus trains, as measured by spike adaptation index. Linear regression line: r^2^ = 0.13, p = 0.015. (**P-V**) Comparisons between bushy neurons with dominant type I_a_ inputs (I_a_-D; I_a_% > 50%) and those with dominant non-type I_a_ inputs (Non-I_a_-D; I_a_% < 50%) in resting potential (**P**), input resistance (**Q**), threshold spike amplitude (**R**), threshold current injection that triggered the first spike (**S**), firing rate throughout the 100 Hz train (**T**), spike adaptation index (**U**), and vector strength of the spikes (**V**). Unpaired t-test or Mann-Whitney test: NS, p > 0.05; *p < 0.05; **p < 0.01; ***p < 0.001. Data are represented as mean ± SD. See also Fig. S1.

### AN synaptopathy is more severe in non-type I_a_ synapses and associated with altered composition of bushy neuron population during ARHL

We next examined the innervation pattern of AN synapses and bushy neuron responses in middle-aged and old mice with different levels of ARHL, as revealed by significantly elevated threshold to clicks in auditory brainstem response (ABR) and reduced ABR wave I amplitude (Fig. 3 A-C). Recordings were obtained from 31 bushy neurons in middle-aged (Fig. 3D-G) and 35 bushy neurons in old (Fig. 3H-K) mice. Similar to the young mice, we observed bushy neurons with both I_a_-D inputs and Non-I_a_-D inputs in middle-aged and old mice (Fig. 3D-K). However, the total volume of VGluT1-labeled puncta per neuron significantly decreased with age (Fig. 3L), suggesting an overall degeneration of AN synapses during ARHL. Interestingly, the degeneration differentially affected type I_a_ (Fig. 3M) and non-type I_a_ (Fig. 3N) synapses, in which it was more profound in non-type I_a_ synapses (Two-way ANOVA: age effect, *F_(2,224)_* = 16.2, *p* < 0.0001; synaptic subtype effect, *F_(1,224)_* = 36.2, *p* < 0.0001). On average, the volume of VGluT1-labeled puncta from type I_a_ synapses decreased by 15% in middle-aged mice and 41% in old mice compared to young mice (Fig. 3M). In contrast, the volume of non-type I_a_ puncta decreased by 34% and 73% in middle-aged and old mice (Fig. 3N), respectively. These results show that there is a progressive AN central synaptopathy during ARHL, and this synaptopathy is biased with more severe degeneration in non-type I_a_ synapses.

**Figure 3.**
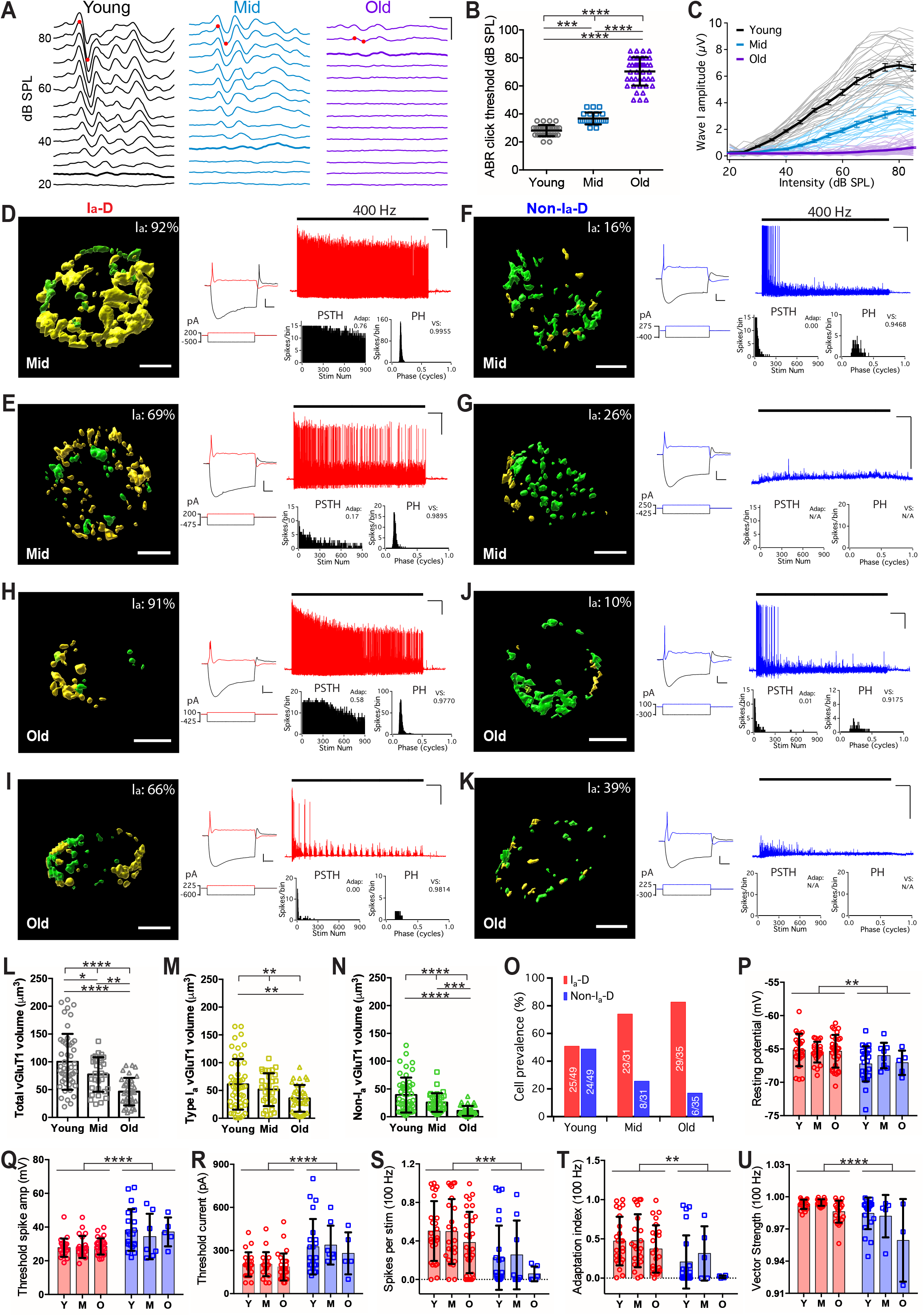
AN synaptopathy is more severe in non-type I_a_ synapses and associated with altered composition of bushy neuron population during ARHL. (**A**) Example ABR waveforms from mice at different ages to clicks at different intensities. Red dots mark the positive and negative peaks of ABR wave I at 80 dB SPL. Thick traces: threshold ABR wave. Scale: 2 ms, 5 μV. (**B**) ABR click thresholds in mice from three age groups. One-way ANOVA and post-hoc tests: ***p < 0.001; ****p < 0.0001. ABR threshold was beyond 85 dB SPL in 12 old mice and not determined. (**C**) Growth curves of ABR wave I amplitude in all mice. Thin lines: individual mice; thick lines: average of each age group, with error bars represent SEM. (**D-K**) Synaptic inputs and response traces of example bushy neurons from middle-aged (**D-G**) and old (**H-K**) mice. Panels are presented in the same pattern as in **Fig. 2A-L**. Left panel: reconstructed VGluT1-stained puncta surrounding the soma of the bushy cell (not shown) from type I_a_ synapses (yellow) and non-type I_a_ synapses (green). I_a_%: volume percentage of the VGluT1-stained puncta from type I_a_ synapses over total. Scale: 5 μm. Middle panel: responses of the bushy cell to current step injections. Scale: 10 mV and 20 ms. Right panels: example response to a train of auditory nerve stimulation (black bar) at 100 Hz. Scale: 10 mV and 1 s. PSTH: post-stimulus time histogram. Adap: spike adaptation index. PH: period histogram. VS: vector strength. **D** is from the same cell in **Fig. 1B-H**. AN stimulation only evoked EPSPs but failed to trigger any spike in **G** and **K**. No adaptation index or vector strength was obtained from these cells. See also Figure S2 for middle aged mice and Figure S3 for old mice. (**L-N**): Comparisons of the total volume (**L**), type I_a_ only volume (**M**) and non-type I_a_ only volume (**N**) of VGluT1-labeled puncta on bushy neurons from three age groups. One-way ANOVA or Kruskal-Wallis with post-hoc tests: *p < 0.05; **p < 0.01; ***p < 0.001; ****p < 0.0001. (**O**) Prevalence of bushy neurons that receive I_a_-D and Non-I_a_-D synaptic inputs among three age groups. Numbers mark the cell count of each type over group total. (**P-U**) Comparisons of bushy neurons among three age groups in resting potential (**P**), threshold spike amplitude (**Q**), threshold current injection that triggered the first spike (**R**), firing rate to 100 Hz stimulation (**S**), spike adaptation index (**T**), and vector strength of the spikes throughout the 100 Hz trains (**U**). Two-way ANOVA revealed significant cell type effect (I_a_-D vs. Non-I_a_-D) in all panels (**P-U**): **p < 0.01; ***p < 0.001; ****p < 0.0001. Age effect was only significant in vector strength (**U**): p < 0.001. Data in **L-N and P-U** are presented as mean ± SD.

Consistent with more profound degeneration of non-type I_a_ synapses, the prevalence of neurons with Non-I_a_-D inputs progressively decreased with age (Fig. 3O). The differences between I_a_-D and non-I_a_-D neurons in intrinsic membrane properties (Fig. 3P-R) and AN-evoked responses (Fig. 3S-U) were retained in middle-aged and old mice (see also Figs. S2 and S3), except that there were more neurons during aging that failed to fire any spike to AN stimulation (Fig. 3G, K) (3/49 in young, 5/31 in middle-aged, 7/35 in old mice). The results suggest an age-related shift in the composition of bushy neuron population during ARHL in the direction of having relatively more I_a_-D neurons.

### Synaptopathy in individual type I_a_ endbulb of Held synapses during ARHL

As calretinin-staining labels the entire type I_a_ synaptic terminal, we reconstructed the 3-dimentional profile of individual type I_a_ endbulb of Held synapses onto bushy neurons from all three age groups of mice (Figs. 1G, 4A-C). On average, bushy neurons with identifiable type I_a_ endbulb terminals received between 1 - 3 type I_a_ endbulbs with an average of 1.5 ± 0.6 (n = 64) in young, 1.5 ± 0.6 (n = 20) in middle-aged, and 1.7 ± 0.7 (n = 32) in old mice (Kruskal-Wallis test: p = 0.472). Individual type I_a_ endbulbs degenerated during aging with significantly decreased volume of the synaptic terminal (Fig. 4D) as well as the volume of VGluT1-labeled puncta within each endbulb (Fig. 4E). However, there was no change in the volume ratio between the two (Fig. 4F), suggesting that the relative quantity of synaptic vesicles remains constant. Thus, AN synaptopathy during ARHL occurs with reduced synaptic terminal volume as well as a balanced decrease in functional components of the synaptic machinery, at least in type I_a_ endbulb of Held synapses.

**Figure 4.**
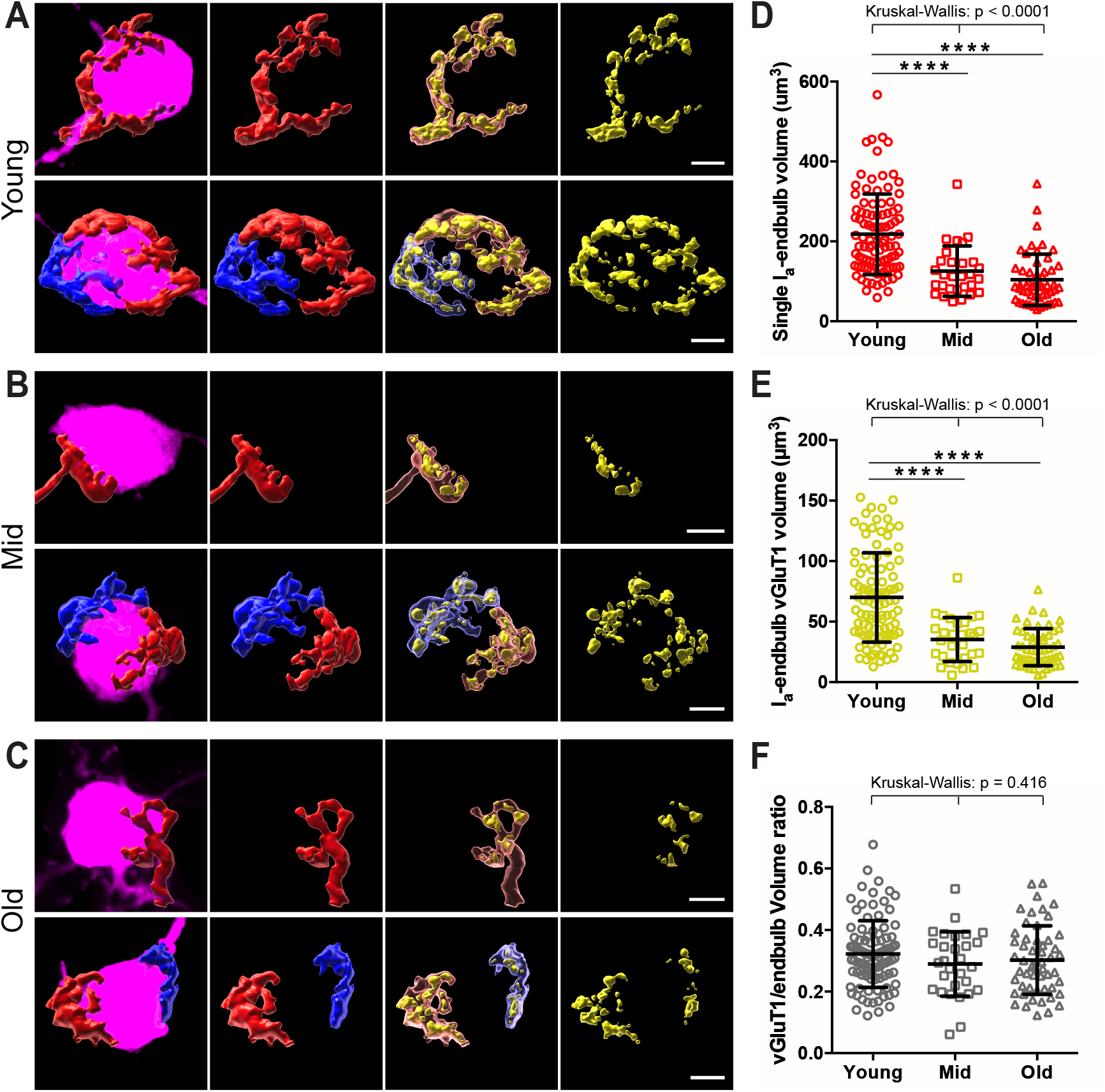
Synaptopathy in individual type I_a_ endbulb of Held synapses during ARHL. (**A**) Morphology of individual type I_a_ endbulb of Held synapses in young mice. Panels from left to right: filled neurons (magenta) with reconstructed type I_a_ endbulbs, type I_a_ endbulbs alone, type I_a_ endbulbs (semi-transparent) with enclosed VGluT1-labled puncta (yellow), and VGluT1-labeled puncta alone. Top panels: example neuron with only one type I_a_ endbulb of Held synapse. Bottom panels: example neuron with two type I_a_ endbulb of Held synapses, which are shown in red and blue respectively. (**B**) Morphology of individual type I_a_ endbulb of Held synapses in middle-aged mice. (**C**) Morphology of individual type I_a_ endbulb of Held synapses in old mice. Scales in **A-C**: 5 μm. (**D-F**) Comparisons of individual type I_a_ endbulb volume (**D**), enclosed VGluT1-puncta volume (**E**), and VGluT1/endbulb volume ratio (**F**) among all three age groups. Kruskal-Wallis and post-hoc tests: ****p < 0.0001. Data are presented as mean ± SD.

## DISCUSSION

### Convergence of different subtypes of AN central synapses

Here we address how different subtypes of SGN neurons project onto individual principal neurons in the CN. The findings provide new insights on how auditory information from the cochlea, especially sound intensity information, is transformed and encoded in CN neurons. Sound intensity at each inner hair cell is conveyed by distributed firing rates amongst three subtypes of type I SGNs that operate over different ranges (Liberman, 1978). How this information is reassembled in CN neurons for central processing has remained unclear despite decades of research. Classical studies relied on tracing individual HRP filled AN fibers with known spontaneous rates (Fekete et al., 1984; Liberman, 1991, 1993; Ryugo, 2008; Ryugo and Sento, 1991; Ryugo et al., 1996; Sento and Ryugo, 1989; Tsuji and Liberman, 1997), but because most of the filled individual fibers ended on different cells, it was not clear how different subtypes of AN synapses converge. Using calretinin as one known marker for type I_a_ SGNs (Petitpre et al., 2018; Shrestha et al., 2018; Sun et al., 2018), we have shown that different subtypes of AN synapses innervate individual CN neurons with a continuous distribution of convergence that correlates with distinct physiological properties of postsynaptic neurons. Our results support the notion that information from different subtypes of SGNs is transformed and processed in the central auditory system in a new dimension that extends even within a single cell type per tranditional classification. Further investigation of this new dimension is essential in understanding the connection between cochlear pathology and central processing disorders under hearing impairments.

### AN synaptopathy and altered composition of bushy neuron population during ARHL

Our findings of a relative loss of type I_b_/I_c_ AN terminals on bushy neurons during ARHL are consistent with age-related selective cochlear synaptopathy in low spontaneous rate/high threshold (type I_c_) SGNs (Furman et al., 2013; Liberman et al., 2015), and suggests that over a long time, type I_b_/I_c_ SGN cell death leads to loss of their central synapses. Such biased AN synaptopathy consequently results in relatively more abundant type I_a_ AN central synapses among the surviving population, which retains more than 70% of all SGNs at 100% life span (Sergeyenko et al., 2013). It remains unclear if the increased prevalence of I_a_-D neurons during ARHL (Fig. 3O) is solely due to the loss of Non-I_a_-D neurons after biased AN synaptopathy, or the conversion of Non-I_a_-D neurons to I_a_-D neurons after adaptive changes like modified expression of voltage-gated ion channels. The observed shift in the composition of neuronal population during ARHL suggests that CN neural circuitry is altered following biased AN synaptopathy, which is expected to impact the central auditory processing in upper nuclei throughout the auditory nervous system.

### Hidden hearing loss in middle-aged mice

CBA/CaJ mice in middle-aged group have relatively normal ABR thresholds but greatly reduced wave I amplitude (Fig. 3A-C), a scenario that mimics hidden hearing loss (Kujawa and Liberman, 2015; Sergeyenko et al., 2013). Both biased AN synaptopathy (Fig. 3L-N) and shifted composition of neuronal population (Fig. 3O) were observed in this age group. The findings indicate that following selective cochlear synaptopathy, which is considered the primary cause of hidden hearing loss, significant changes in central structures along the auditory pathway also occur at this stage before hearing loss becomes substantial. In particular, bushy neurons are specialized in processing temporal information (Joris et al., 1994a; Joris et al., 1994b). Decreased prevalence of Non-I_a_-D bushy neurons (Fig. 3O) in middle aged mice suggests that the temporal processing of high intensity information from non-type I_a_ SGNs is compromised in CN neural circuitry under hidden hearing loss. Since CN neurons initiate parallel pathways in the central auditory system (Cant and Benson, 2003), the shifted composition of CN neuronal population could be the trigger of many central auditory processing disorders including hyperacusis (loudness intolerance) and tinnitus (Auerbach et al., 2014; Caspary and Llano, 2017; Luo et al., 2017; Niu et al., 2013; Radziwon et al., 2019; Sheppard et al., 2019; Shore and Wu, 2019; Vogler et al., 2011; Wang et al., 2011).

### Clinical implications

Cochlear implantation has been widely used in the clinic to treat hearing loss. Its performance relies on the function of SGNs. Age-related AN central synaptopathy is no doubt one of the factors that limit the performance of cochlear implants. Indeed, duration of deafness is the most important predictor of the postimplant outcomes (Rubinstein et al., 1999). The stimulation parameters of modern cochlear implants activate all SGNs indiscriminately. Our study suggests that, if possible, differential stimulation of different subtypes of SGNs in cochlear implants with enhanced activation of non-type I_a_ SGNs may compensate the biased AN central synaptopathy and altered CN neuronal census, which would enhance auditory processing, improve cochlear implant performance and overall hearing.

## ACKNOWLEDGMENTS

We thank Paul B. Manis for comments on the manuscript. This work was supported by NIH grant R01DC016037 to R.X.. Images and video were generated at OSU Campus Microscopy and Imaging Facility, supported by grant P30CA016058.

## AUTHOR CONTRIBUTIONS

R.X. designed the research; R.X., M.W., C.Z., S.L., and Y.W. conducted the research; M.W., R.X., B.J.S., and R.W.A. analyzed the data; R.X. wrote the paper; All authors edited the paper.

## STAR METHODS

### RESOURCE AVAILABILITY

#### Lead Contact

Further information and requests for resources and reagents should be directed to and will be fulfilled by the Lead Contact, Ruili Xie.

#### Materials Availability

This study did not generate new unique reagents.

#### Data and Code Availability

The article includes all data generated during this study. Custom-written procedures in Igor Pro were used in the analysis of all electrophysiological data and will be available upon request.

### EXPERIMENTAL MODEL AND SUBJECT DETAILS

CBA/CaJ mice were purchased from The Jackson Laboratory, bred and maintained at the animal facility at The Ohio State University. Experiments used 32 young mice (1.5-4.5 months), 22 middle-aged mice (17-19 months), and 55 aged mice (28-32 months) of either sex. All experiments were conducted under the guidelines of the protocols approved by the Institutional Animal Care and Use Committees at The Ohio State University.

### METHOD DETAILS

#### Auditory Brainstem Response (ABR)

Hearing status of the mice were assessed by measuring ABR to clicks as previously described (Wang et al., 2019; Xie, 2016). Briefly, mice were anaesthetized with IP injection of ketamine (100 mg/kg) and xylazine (10 mg/kg) and placed inside a sound-attenuating chamber. Body temperature was maintained at ~ 36 °C using a feedback-controlled heating pad. ABR to clicks were acquired using a RZ6-A-P1 system with BioSigRZ software (Tucker-Davis Technologies). Clicks (0.1 ms, monophasic with alternating phase; 21 times/s) were delivered through a free field MF1 magnetic speaker 10 cm away from the pinna. Recording needle electrodes were placed at the ipsilateral pinna and vertex, with the ground electrode at the rump. ABR wave at each sound level was averaged from 512 repetitions.

#### Brain Slice Preparation

Under ketamine/xylazine anesthesia, mice were decapitated and the skulls were opened to retrieve the brainstem. Parasagittal slices containing the cochlear nucleus were cut at a thickness of 225 - 240 μm using a Vibratome 1000 (Technical Products, Inc.) or a VT1200S Microtome (Leica Biosystems). Slices were then incubated in artificial cerebral spinal fluid (ACSF) at 34 °C for 30-45 minutes before recordings began. ACSF contained (in mM): 122 NaCl, 3 KCl, 1.25 NaH_2_PO_4_, 25 NaHCO_3_, 20 glucose, 3 *myo*-inositol, 2 sodium pyruvate, 0.4 ascorbic acid, 1.8 CaCl_2_ and 1.5 MgSO_4_, and was constantly gassed with 95% O_2_ and 5% CO_2_.

#### Electrophysiological Recording

After incubation, the brain slice was moved to an ACSF-bathed recording chamber under an Axio Examiner microscope (Carl Zeiss). Whole-cell recordings in current clamp mode were performed from bushy neurons of the AVCN. Recording pipettes were made using a P-2000 micropipette puller (Sutter Instrument) and filled with electrode solution that contained (in mM): 126 potassium gluconate, 6 KCl, 2 NaCl, 10 HEPES, 0.2 EGTA, 4 MgATP, 0.3 GTP, 10 Tris-phosphocreatine, and pH adjusted to 7.20. Alexa Fluor 594 was added to the electrode solution at the final concentration of 0.01% by weight to dye-fill the neuron for online visualization of cell morphology as well as cell labeling upon completion. Data acquisition used hardware and software from Molecular devices, including Multiclamp 700B amplifier, Digidata 1550B acquisition system and pClamp 11 software. All recordings were made at 34 °C. Only bushy neurons were included in the study, which were identified based on electrophysiological response properties and morphological features as previously described (Cant and Morest, 1979; Manis et al., 2019; Webster and Trune, 1982; Wu and Oertel, 1984). Responses to current step injections were obtained to assess intrinsic membrane properties. Auditory nerve stimulation (Fig. 1A) was delivered through a 75 μm diameter concentric stimulating electrode (Frederick Haer Company). The stimulus pulse had a duration of 0.1 ms, with intensity set at ~30% above the threshold intensity level that first triggered spikes in the target neuron. For neurons that failed to fire any spikes, stimulus intensity was tested up to the level that would cause tissue damage at the stimulation site. Responses to trains of auditory nerve stimulation at 100 and 400 Hz were recorded. After completion of data acquisition, the recording electrode was slowly withdrawn until it broke off from the target neuron, which usually resealed itself with preserved neuronal morphology and remained filled with Alexa Fluor 594 dye. The brain slice was immediately fixed in 4% paraformaldehyde in PBS for 15 minutes, followed by post-hoc immunostaining.

#### Immunohistochemistry

Fixed brain slices with dye-filled neurons were processed for immunostaining as previously described (Karadottir and Attwell, 2006; Lin and Xie, 2019). After rinsing in PBS (3 times, 15 minutes each), slices were pre-incubated in blocking solution (10% horse serum, 0.5% Triton X-100, and 0.05% NaN_3_ in PBS) for 6 hours at room temperature, followed by overnight incubation with primary antibodies against VGluT1 (guinea pig anti VGluT1; Cat# 135304, Synaptic Systems, 1:500) and calretinin (rabbit anti calretinin; Cat# 214102, Synaptic Systems, 1:500) at 4 °C. Slices were then rinsed again in PBS (4 times, 20 minutes each), incubated with secondary antibodies (goat anti guinea pig IgG, Alexa 488 conjugated; Cat # A11073, Thermo Fisher Scientific, 1:500; and goat anti rabbit IgG, Alexa 647 conjugated; Cat# A21245, Thermo Fisher Scientific, 1:500) at room temperature for 4 hours, re-rinsed, and mounted on slide with DAPI-Fluoromount-G mounting media (Southern Biotech). Filled neurons and labeled auditory nerve synapses were imaged using an Olympus FV3000 confocal microscope. Images were sampled using a 60x oil immersion objective, 3.0 times software zoom, a z-step size of 0.3 μm, and at a resolution of either 800 × 800 or 1024 × 1024 pixels.

#### Image processing

All image processing and measurement were done using Imaris software (version 9.5.0; Oxford Instruments). Three-dimensional reconstruction of the dye-filled target bushy neuron (magenta), VGluT1-labeled puncta (green) and type I_a_ endbulb of Held terminals (red) in close proximity to the soma of the target neuron were made by generating 3D surfaces in three separate channels using the semi-automatic method. VGluT1-labeled puncta that located inside type I_a_ endbulbs were determined as puncta from type I_a_ synapses and marked in yellow, while those outside of type I_a_ endbulbs were from non-type I_a_ synapses and marked in green (Fig. 1D-F, Supplemental Video 1). Volumes of VGluT1-labeled puncta and individual recognizable endbulb of Held synapses were measured.

#### Electrophysiology Analysis

Electrophysiological data were analyzed in Igor Pro (WaveMetrics) using custom-written functions. Responses to current step injections were used to quantify the intrinsic membrane properties as previously described (Xie, 2016; Xie and Manis, 2017a). Input resistance was calculated as the slope of the current voltage relationship curve between current injection of −25, −50, −75, and −100 pA and the resulted peak hyperpolarization amplitudes. Depolarizing current step injection was gradually increased by the step of 25 pA until the target neuron fired the first spike, i.e. the threshold spike. This current level was determined as the threshold current of the neuron. Threshold spike amplitude was calculated as the voltage difference between the resting membrane potential and the peak of the threshold spike. Spike adaptation index of the responses to 100 and 400 Hz trains were calculated as the ratio of the firing rate during the last 5 seconds of the train to the firing rate during the first second. Vector strength (Goldberg and Brown, 1969) of the stimulus train-evoked spikes was calculated using the formula: 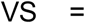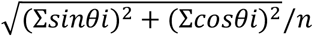, where θ_*i*_ is the phase angle of the spike *i* relative to the stimulation cycle, and n is the total number of spikes throughout the stimulus trains.

#### Statistical Analysis

Statistical analysis used GraphPad Prism (version 6.0h). Kolmogorov-Smirnov test was first performed to check if population data are normally distributed. As stated in the text, statistical comparisons of group data with normal distribution were made using unpaired t-test or one-way ANOVA followed by Tukey’s multiple comparisons test. Comparisons of group data that are not normally distributed used Mann Whitney test or Kruskal-Wallis test followed by Dunn’s multiple comparisons test. Two-way ANOVA was also used (Fig. 3P-U) as stated. All data are presented as mean ± SD except in Fig. 3C, which plotted mean ± SEM for clarity purpose of the panel.

## SUPPLEMENTAL INFORMATION

**Supplemental Video 1.** Three-dimensional reconstruction of the target neuron and connected synapses. Background music: *The Story Unfolds* - Jingle Punks (https://youtu.be/_8iypSHvdx0)

**Figure S1.**
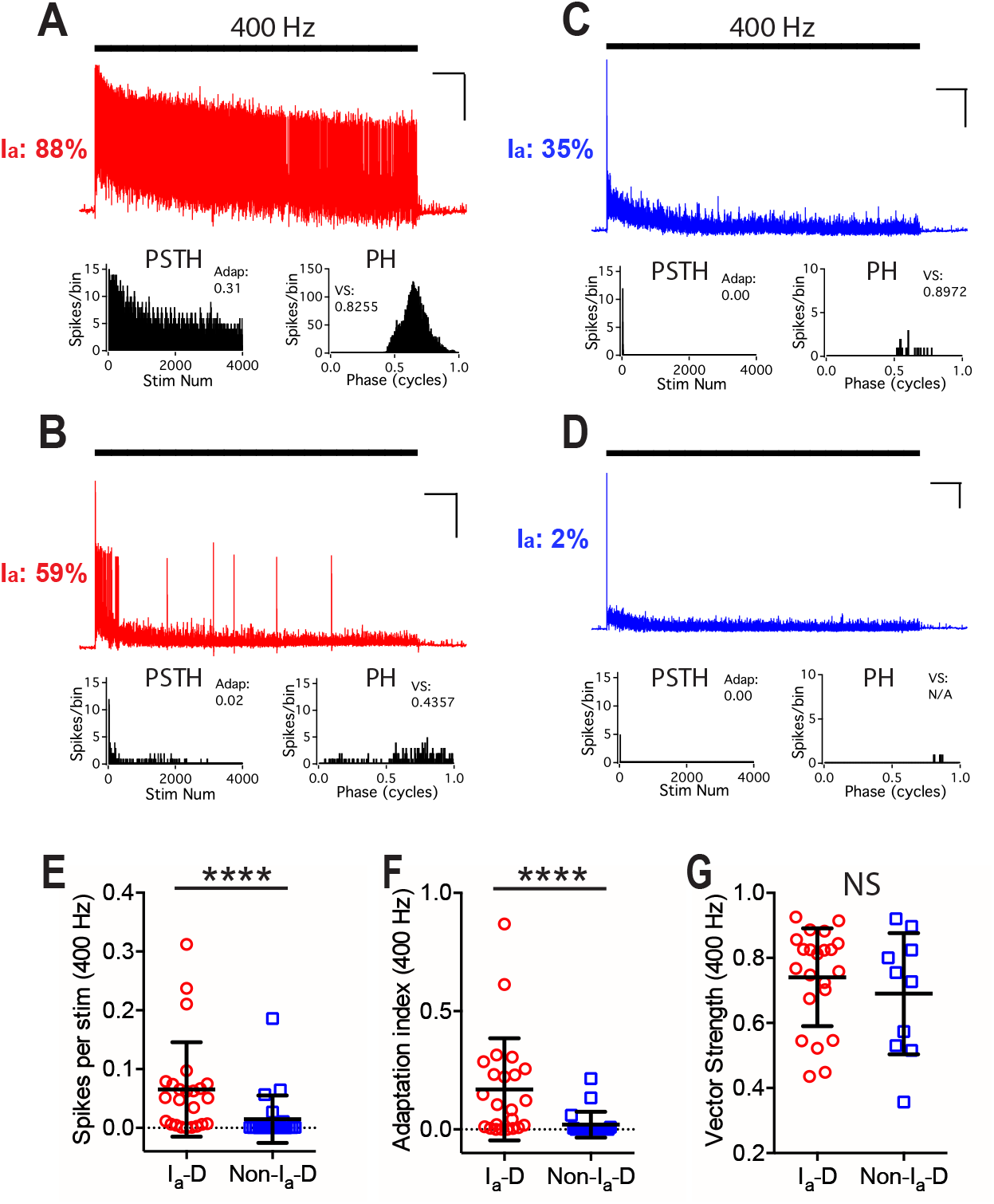
Bushy neurons with different convergence of AN synaptic inputs fire differently to trains of AN stimulation at 400 Hz. (**A**) The same example bushy neuron in Fig. **2A** fired sustained spikes to trains of AN stimulation at 400 Hz. This neuron received mostly type I_a_ synaptic inputs (88% in I_a_-VGluT1 puncta volume as shown in Fig. **2A**). Scale: 10 mV and 1 s. PSTH: post-stimulus time histogram. Adap: spike adaptation index. PH: period histogram. VS: vector strength. (**B-D**) Responses of bushy neurons in Fig. **2D**, **G**, and **J** to AN stimulation at 400 Hz. Vector strength of the cell in **D** was not calculated due to insufficient number of spikes. (**E-G**) Comparisons between bushy neurons with dominant type I_a_ inputs (I_a_-D; I_a_% > 50%) and those with dominant non-type I_a_ inputs (Non-I_a_-D; I_a_% < 50%) to 400 Hz AN stimulus trains in firing rate (**E**), spike adaptation index (**F**), and vector strength (**G**). Vector strength in 11 out of 21 neurons that received Non-I_a_-D inputs (blue) was not available due to insufficient number of spikes. Unpaired t-test or Mann Whitney test: NS, p > 0.05; ****p < 0.0001.

**Figure S2.**
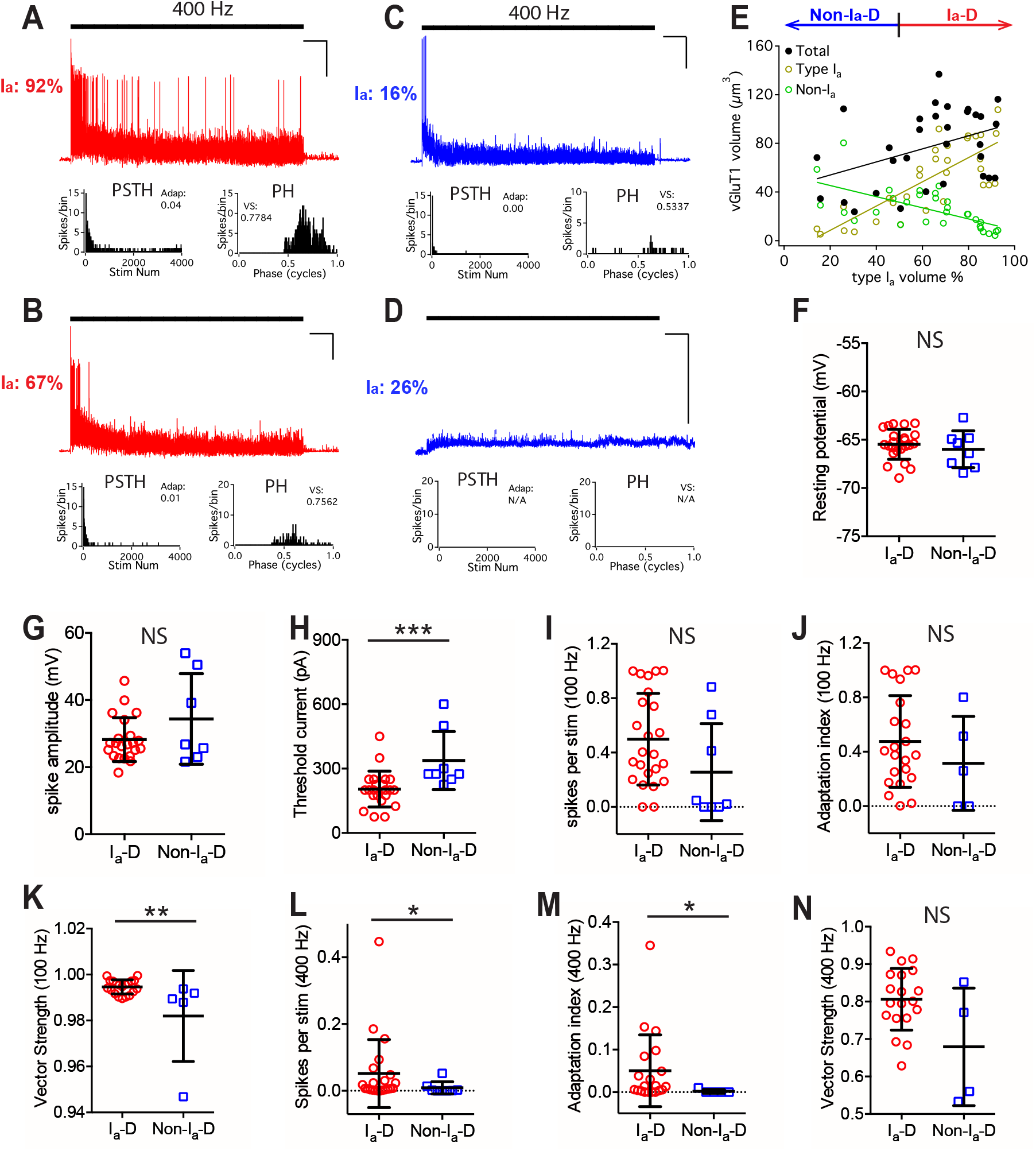
Properties of bushy neurons from middle-aged mice with different convergence of AN synaptic inputs. (**A-D**) Responses of the same example bushy neurons in Fig. **3D-G** to trains of AN stimulation at 400 Hz. Cells in red are I_a_-D neurons that receive over 50% of type I_a_-puncta volume. Cells in blue are Non-I_a_-D neurons with less than 50% of type I_a_-puncta volume. Scale: 10 mV and 1 s. PSTH: post-stimulus time histogram. Adap: spike adaptation index. PH: period histogram. VS: vector strength. No spike was evoked from the cell in **D**, and no adaptation index or vector strength was calculated. (**E**) Bushy neurons with different fraction of type I_a_ inputs (x-axis) show correlated distribution in their total volume of VGluT1-stained puncta (black), volume of type I_a_ only puncta (yellow), and volume of non-type I_a_ only puncta (green). Linear regression lines: black, r^2^ = 0.16, p = 0.025; yellow, r^2^ = 0.62, p < 0.0001; green, r^2^ = 0.40, p < 0.0001. (**F-H**) Comparisons between the intrinsic property of I_a_-D and Non-I_a_-D bushy neurons from the middle-aged mice, including threshold current (**F**), threshold spike amplitude (**G**), and threshold current injection that evoked first spikes (**H**). (**I-N**) Comparisons of firing properties between I_a_-D and Non-I_a_-D bushy neurons in response to 100 Hz (**I-K**) and 400 Hz trains (**L-N**), including firing rate, spike adaptation index and vector strength. Data are represented as mean ± SD. Unpaired t-test or Mann-Whitney test: NS, p > 0.05; *p < 0.05; **p < 0.01; ***p < 0.001.

**Figure S3.**
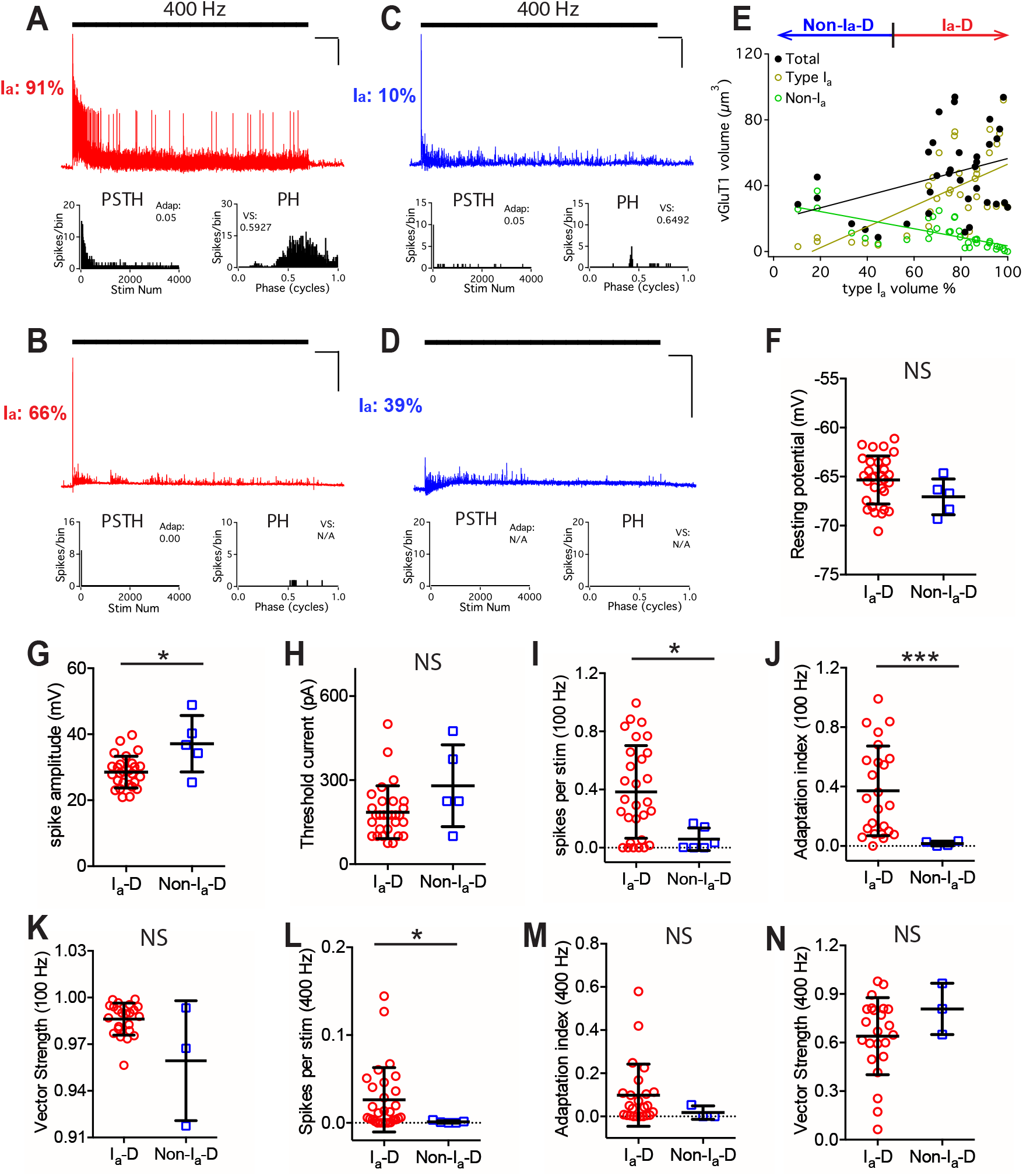
Properties of bushy neurons from old mice with different convergence of AN synaptic inputs. (**A-D**) Responses of the same example bushy neurons in Fig. **3H-K** to trains of AN stimulation at 400 Hz. Cells in red are I_a_-dominant neurons that receive over 50% of type I_a_-puncta volume. Cells in blue are non-I_a_-dominant neurons with less than 50% of type I_a_-puncta volume. Scale: 10 mV and 1 s. PSTH: post-stimulus time histogram. Adap: spike adaptation index. PH: period histogram. VS: vector strength. No spike was evoked from the cell in D, and no adaptation index or vector strength was calculated. (**E**) Bushy neurons with different fraction of type I_a_ inputs (x-axis) show correlated distribution in their total volume of VGluT1-stained puncta (black), volume of type I_a_ only puncta (yellow), and volume of non-type I_a_ only puncta (green). Linear regression lines: black, r^2^ = 0.13, p = 0.033; yellow, r^2^ = 0.40, p < 0.0001; green, r^2^ = 0.49, p < 0.0001. (**F-H**) Comparisons between the intrinsic property of I_a_-D and Non-I_a_-D bushy neurons from old mice, including resting potential (**F**), threshold spike amplitude (**G**), and threshold current injection that evoked first spikes (**H**). (**I-N**) Comparisons of firing properties between I_a_-D and Non-I_a_-D bushy neurons in response to 100 Hz (**I-K**) and 400 Hz trains (**L-N**), including firing rate, spike adaptation index and vector strength. Data are represented as mean ± SD. Unpaired t-test or Mann-Whitney test: NS, p > 0.05; *p < 0.05; ***p < 0.001.

---

KEY RESOURCES TABLE.

KEY RESOURCES TABLE
| REAGENT or RESOURCE | SOURCE | IDENTIFIER |
| --- | --- | --- |
| **Antibodies** |  |  |
| Guinea pig anti-vGluT1 | Synaptic Systems | RRID: AB_887878 |
| Rabbit anti-calretinin | Synaptic Systems | RRID: AB_2228331 |
| Goat anti-guinea pig, Alexa 488 conjugated | Thermo Fisher Scientific | RRID: AB_2534117 |
| Goat anti-rabbit, Alexa 647 conjugated | Thermo Fisher Scientific | RRIP: AB_2535813 |
| **Bacterial and Virus Strains** |  |  |
| **Biological Samples** |  |  |
| **Chemicals, Peptides, and Recombinant Proteins** |  |  |
| **Critical Commercial Assays** |  |  |
| **Deposited Data** |  |  |
| **Experimental Models: Cell Lines** |  |  |
| **Experimental Models: Organisms/Strains** |  |  |
| Mouse: CBA/CaJ | Jackson Laboratory | Stock #: 000654 |
| **Oligonucleotides** |  |  |
| **Recombinant DNA** |  |  |
| **Software and Algorithms** |  |  |
| Igor Pro v 6.37 | WaveMetrics | https://www.wavemetrics.com |
| Imaris v | Oxford Instruments | https://imaris.oxinst.com |
| Prism v 6.0h | GraphPad | https://www.graphpad.com/scientific-software/prism/ |
| **Other** |  |  |

